# A reduced-order multibody model of the foot–ankle complex based on kinematic synergies

**DOI:** 10.64898/2026.05.17.725725

**Authors:** Michele Conconi, Luca Modenese, Gian Marco Barbieri, Alberto Leardini, Claudio Belvedere, Nicola Sancisi

## Abstract

**Background and Objective:** The foot–ankle complex is a highly articulated and mechanically constrained system, often simplified as a chain of few rigid segments, neglecting many bone-to-bone motions and raising questions about the accurate representation of interaction with ground. This study proposes a new reduced-order multibody formulation that captures intrinsic kinematic constraints of the foot through motion synergies.

**Methods:** Bones kinematic coupling, or motion synergies, were experimentally derived from weight-bearing CT scans using principal component analysis. These couplings were embedded in a synergy-based multibody kinematic optimization framework describing the foot–ankle with five degrees of freedom: ankle flexion; foot adduction, pronation, and arching; and toe flexion. Model accuracy was evaluated against bone-level experimental kinematics. The model was applied to gait data from healthy, flat, and diabetic feet and compared with a standard multi-segment foot model, assessing robustness by progressively reducing the number of skin markers.

**Results:** Average errors were about 1° and 0.5 mm when using subject-specific synergies and below 7° and 4 mm when scaling the generic model, matching or exceeding the accuracy of existing models. Reliable reconstruction was obtained using only four foot markers. In clinical gait analysis, the model showed superior discrimination between populations and enabled assessment of transverse arch deformation, not accessible with conventional models.

**Conclusion:** The proposed synergy-based model provides an accurate, low-complexity framework for reconstructing bone-level foot and ankle kinematics, substantially simplifying gait analysis while improving biomechanical interpretability. This framework supports future integration with dynamic models aimed at studying load transmission in the foot.

## 1 INTRODUCTION

The foot–ankle complex poses a major challenge for biomechanical modeling due to its high level of articulation, with the motion of 26 small bones contributing to overall function. To manage this complexity, most gait analysis approaches group foot bones into clusters assumed to behave as rigid segments, whose motion is tracked using skin-mounted markers. This strategy, commonly referred to as multi-segment foot modeling (MSFM), has been widely adopted over the last decades. However, no consensus exists regarding the optimal clustering strategy, with proposed MSFMs ranging from 3 to 11 segments (Leardini, Caravaggi, Theologis, & Stebbins, 2019). Comparisons with more accurate yet invasive quantification of foot kinematics, such as bone pins (Nester, et al., 2007) (Lundgren, et al., 2008) (Le, et al., 2025) or biplanar fluoroscopy (Ito, et al., 2015) (Maharaj, et al., 2020) have highlighted the inability of MSFM approaches to capture the details of the relative bone-to-bone kinematics.

These limitations arise partly from marker-based motion capture, which is affected by soft tissue (STA) (Leardini, Chiari, Della Croce, & Cappozzo, 2005) (Kessler, et al., 2019) and marker occlusion artefact (MOA) (Conconi, Pompili, Sancisi, & Parenti-Castelli, Quantification of the errors associated with marker occlusion in stereophotogrammetric systems and implications on gait analysis, 2021), and partly from the modeling assumptions themselves. In particular, clustering inherently neglects relative motion among bones within each segment, while direct kinematic formulations may lead to non-physiological joint disarticulation and bone interpenetration.

To address these issues, multibody kinematic optimization (MKO) frameworks have been proposed (Lu & O’connor, 1999) (Leardini, et al., 2017), in which a kinematic chain constrains the relative motion of the skeletal segments, and optimal marker tracking is achieved by fitting the entire chain to the experimental data. Existing MKO-based foot models vary widely in the number of segments (3 to 24), joint definitions, degrees of freedom (DOFs), and the presence of implicit couplings among DOFs (Saraswat, Andersen, & MacWilliams, 2010) (Scott & Winter, 1993) (Montefiori, et al., 2019) (Malaquias, et al., 2017) (Oosterwaal, et al., 2016). Notably, increasing model complexity does not necessarily improve biomechanical fidelity: higher-DOF models have been shown to reduce the ability to reproduce physiological foot kinematics (Gholami, Pàmies-Vilà, Kövecses, & Font-Llagunes, 2015) (Malaquias, et al., 2017), while overly simplified serial-chain models neglect mediolateral load transfer, leading to inaccurate kinetic estimates(Buczek, Walker, Rainbow, Cooney, & Sanders, 2006). Moreover, increasing the number of DOFs substantially raises experimental complexity, as more markers are required to reconstruct kinematics, amplifying STA and MOA effects and reducing repeatability and comparability across measurements (Bruening, Cooney, & Buczek, Analysis of a kinetic multi-segment foot model. Part I: Model repeatability and kinematic validity, 2012) (Carson, Harrington, Thompson, O’connor, & Theologis, 2001). Likely for these reasons, early full-body musculoskeletal models often represented the foot as two rigid segments articulated at the toes (Damsgaard, Rasmussen, Christensen, Surma, & De Zee, 2006) (Delp, et al., 1990) (Dhang, Rasmussen, Damsgaard, Christensen, & de Zee, 2004) (Hoy, Zajac, & Gordon, 1990) (Saraswat, Andersen, & MacWilliams, 2010), neglecting intrinsic foot motion and typically overestimating ankle joint power by attributing midfoot motion to the ankle joint (Bruening, Cooney, & Buczek, Analysis of a kinetic multi-segment foot model part II: kinetics and clinical implications, 2012).

Together, these considerations highlight the need for a foot model capable of representing bone-level kinematics with a reduced number of DOFs, while preserving mechanical consistency and minimizing experimental burden. One strategy is to explicitly model ligamentous and articular constraints; however, fully explicit formulations require extensive parameter tuning and entail high computational costs (Sikidar & Kalyanasundaram, 2022).

An alternative approach is to represent articular and ligamentous constraints implicitly, rather than modeling them explicitly. In a recent study, the present authors demonstrated that foot–ankle kinematics can be effectively described as a four-degree-of-freedom system using motion synergies, defined as principal kinematic couplings among the rotations and translations of all bones in the foot complex (Conconi, Sancisi, Leardini, & Belvedere, 2024). Related concepts, referred to as *functional units* or *rhythms*, had been previously introduced based on measurements from intracortical pins placed in a limited set of foot bones, including the calcaneus, talus, cuboid, navicular, medial cuneiform, and selected metatarsals (Wolf, et al., 2008) (Oosterwaal, et al., 2016). Due to experimental challenges in accurately locating tracking devices relative to the bones and to the limited number of instrumented segments, these earlier approaches provided primarily qualitative descriptions of kinematic coupling, restricted to subsets of the foot articulations.

In contrast, the work of (Conconi, Sancisi, Leardini, & Belvedere, 2024) reconstructed the three-dimensional kinematics of all foot bones from a series of weight-bearing CT scans, adopting anatomical reference systems (ARS) specifically designed to minimize secondary motion components (Conconi, et al., New anatomical reference systems for the bones of the foot and ankle complex: definitions and exploitation on clinical conditions, 2021). This formulation enabled accurate and consistent tracking of bone positions and orientations across postures and subjects and, through principal component analysis (PCA), led to the identification of one dominant synergy at the ankle and three synergies within the foot. Together, these synergies accounted for more than 95% of the overall mobility of the foot–ankle complex. Importantly, the identified synergies were highly correlated across subjects and were associated with mechanically and clinically meaningful modes of motion, including ankle flexion, Chopart joint abduction, foot pronation, and arch deformation during load acceptance (Conconi, Sancisi, Leardini, & Belvedere, 2024).

Building on these findings, the present study introduces a synergy-based multibody kinematic optimization (SB-MKO) model that enables reconstruction of the kinematics of all 26 foot bones using only five DOFs: the four foot and ankle synergies identified in (Conconi, Sancisi, Leardini, & Belvedere, 2024) plus an additional synergy accounting for toe flexion. Model accuracy and robustness are first evaluated when reconstructing bone motion using in-vitro data, also examining the impact of reducing the number of markers. The model is then applied to gait data from three clinical populations and compare its ability to discriminate between healthy and pathological feet against the standard Rizzoli MSFM model. The proposed model is implemented and made freely available in both MATLAB and OpenSim (Conconi, 2026) (Modenese, 2026).2

## MATERIAL AND METHODS

### 2.1 Experimental quantification of motion synergies

Following the protocol described in (Conconi, Sancisi, Leardini, & Belvedere, 2024), we analyzed six fresh-frozen full lower limbs from cadaveric donors with no signs of previous pathology (three males, three females; age: 70.5 ± 5.9 years; height: 171.5 ± 5.2 cm; weight: 95.8 ± 13.6 kg). The three female donors were analyzed for the first time in this study.

Each specimen was scanned using a CBCT system (‘OnSight 3D Extremity System’, Carestream, Rochester, NY-USA; scan rotation angle 215°; volumetric data: 884×884×960, isotropic voxels of 0.26 mm; tube voltage and current set at 90 kVp and 5.0 mA). The leg was positioned vertically in the scanner, while ankle dorsi/plantarflexion was adjusted to five angles (−30°, −10°, 0°, 10°, 20°) and foot pronation/supination to three angles (−10°, 0°, 10°), yielding 15 unique orientations using wooden wedges. These 15 postures were imaged both unloaded and under weightbearing. Two additional scans were acquired with the leg internally and externally rotated. In total, 32 CBCT scans were obtained for each leg (15 unloaded and 17 loaded). Further methodological details are available in (Conconi, et al., Foot kinematics as a function of ground orientation and weightbearing, 2023).

For each scan, 3D bone models of the 14 bones of the foot and ankle (toes excluded) were reconstructed through a specific automatic software (Disior^®^ v2.1.5 Bonelogic− (SMART28), Disior Ltd. - in collaboration with Paragon 28, Inc. and Zimmer Biomet - Helsinki, Finland). Anatomical reference systems were defined for each bone in the neutral scan (Conconi, et al., New anatomical reference systems for the bones of the foot and ankle complex: definitions and exploitation on clinical conditions, 2021). The neutral STL model of each bone was then registered to all other scans using an automatic ICP process implemented in MATLAB, determining the rototranslational matrix that describes the pose of each bone in each scan. The global foot posture ***p*** was then defined in terms of the relative positions and orientations of the bones. Specifically, the motion of talus (TA) and fibula (FI) were expressed relative to the tibia (TI) absolute pose; the motion of the calcaneus (CA) and the navicular (NA) relative to TA; the motion of the remaining bones (CU, cuboid; CM, medial cuneiform; CI, intermediate cuneiform; CL, lateral cuneiform; M1–5, first to fifth metatarsals) relative to NA. This strategy reduces the number of transformations required to reconstruct the global foot posture ***p*** and results in a comparable range of motion across joints. The posture vector ***p*** contains the concatenated positional and orientational coordinates of all the bones, so that

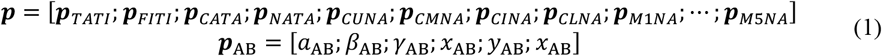

where ***p***_AB_ represent the pose of the proximal bone A in the reference system of the distal bone B, α is the rotation about the x axis of the distal bone (mapping the pronation/supination), γ is the rotation about the z axis of the proximal bone (mapping the dorsi/plantar flexion), and β is the rotation about the floating axis, i.e. perpendicular to the previous two (mapping abduction/adduction), while x, y, and z represent the coordinates of the system origin. A more detailed description of this parametrization can be found in the supplementary material.

Following the procedure described in (Conconi, Sancisi, Leardini, & Belvedere, 2024), PCA was performed on the dataset of *n*=32 postures collected for each foot, separating the ankle (TI, TA, FI) from the rest of the foot. Without loss of generality and following the notation in (Moore, Kooijman, Schwab, & Hubbard, 2011), the mean posture vector is computed at first as

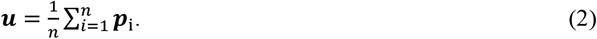

We then computed the centered posture matrix and the corresponding covariance as

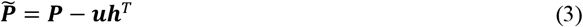

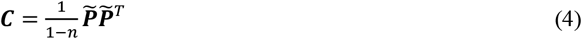

where ***P*** = [***p***_1_, … , ***p***_n_], and ***h*** is an nx1 vector of ones. Principal components or synergies ***s***_*i*_ can be obtained by computing the eigenvectors of ***C***. The generic foot posture can be reconstructed using a linear combination of synergies as

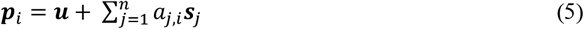

where the ***a*** coefficients can be found by solving the linear system

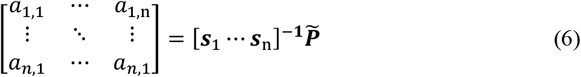

### 2.2 Construction of the synergy-based model

To develop a simplified but representative synergy-based model, we identified the minimal subset of synergies capable of reconstructing foot postures with minimal information loss while remaining consistent across specimens. The Eckart-Young-Mirsky theorem guarantees that the optimal approximation of the reduced problem can be obtained by considering the synergies to which the highest eigenvalues of ***C*** are associated.

The optimal number of synergies in this subset can be obtained through the analysis of the cumulated variance: for all the feet considered in this study, one single synergy explains more than 95% of total ankle mobility, and three synergies are enough to explain more than 92.5% of the rest of the foot mobility (Fig. S1). To verify whether these synergies are also common across feet, we computed inter-subject correlations among corresponding synergies from different subjects (Table 1). Results confirmed previous findings (Conconi, Sancisi, Leardini, & Belvedere, 2024), suggesting that the first three synergies, mapping the dorsi-plantar flexion, the foot abduction/adduction at the Chopart and subtalar joint, and the overall foot pronation/supination, respectively, are indeed invariant among the feet. The fourth foot synergy, mapping the opening of the foot arches during load acceptance, showed lower correlation, suggesting greater variability in foot deformation under loads. A visual representation can be found in the supplementary material (Figure S2 and S3).

**Table 1:**
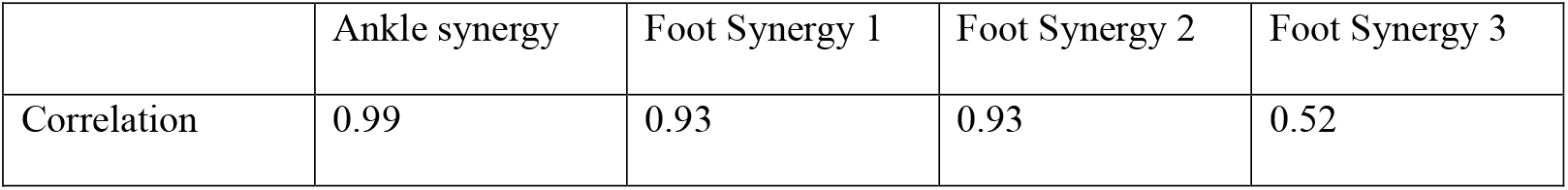
Inter-subject correlation among corresponding synergies from the six specimens analyzed in this study.

Based on these observations, an average foot model was defined. The specimen with the foot length closest to the population mean was selected as reference, where foot length was defined as the distance between the most posterior calcaneal point and the most anterior second-metatarsal-head point in the neutral unloaded posture. The mean posture ***u*** and the four main synergies ***s***_*j*_ of each subject (including the reference one) were normalized according to foot-length ratio by isotropically scaling the translational components. Finally, the average foot mean posture and synergies were computed as the mean of the normalized values. The resulting numerical data defining the average model are provided online at (Conconi, 2026).

To complete the kinematic structure, the toes were modeled as five rigid bodies, each connected to the corresponding metatarsal via a hinge joint whose axis was determined by fitting a cylinder to the metatarsal head articular surface. Toe flexion was coupled through an additional synergy linking the rotations of all toes to the rotation of the hallux.

To complete the model, a set of virtual markers was manually added to the foot bones of the reference subject, in the neutral posture, following the Rizzoli protocol (Leardini, et al., 2007). The resulting average model was mirrored to generate left and right versions.

For MKO, the generic foot model is anisotropically scaled to each subject using a static trial, as described in the Supplementary Material. The motion of the tibia and the foot is then reconstructed by minimizing, frame by frame, the error between the experimental and model markers, whose locations are determined using 11 DOF: 6 of them locate the shank in space, while the remaining 5 are the ankle, foot, and toes synergy coefficients. The detailed computations are again reported in the supplementary material and in the MATLAB code associated with the paper (Conconi, 2026).

### 2.3 OpenSim Implementation

A version of the synergy-based foot model was developed in OpenSim 4.5 (Seth, et al., 2018) (Delp, et al., 2007). A total of 19 rigid bodies, corresponding to the bones of the foot model, with each toe treated as a single body, were created. These bodies were then connected by 19 joints (*CustomJoint* in OpenSim), each presenting 6 DOF. Except for the ground-tibia joint, for which three tibial rotations and translations with respect to the ground were maintained, the other joints’ coordinates were coupled through the synergy coefficients using the linear relationships from the experimental findings. The couplings were implemented in the model via constraints (*CoordinateCouplerConstraints*), solving for the root of multivariate polynomial functions in which the synergy coefficients were the independent variables and the joint coordinates the dependent ones.

The synergy coefficients, which, despite having a functional meaning, are not associated with a physical joint coordinate, were explicitly introduced in the model as the translational degrees of freedom of auxiliary bodies of negligible inertial properties. With this setup, the standard inverse kinematic tool available in OpenSim could be employed to compute foot kinematics from experimental marker data, solving for the tibial coordinates and synergy coefficients as independent coordinates. All inverse kinematic analysis were performed with accuracy 1e-9, instead of the default 1e-5, to ensure numerical robustness. The joint kinematics computed through the OpenSim average model were verified to be identical to those yielded by its MATLAB implementation.

Following a custom scaling of the model implemented in MATLAB, patient specific OpenSim foot models can be created automatically through a script leveraging the software API (Modenese, 2026).

### 2.4 Accuracy of synergy-based description of the foot posture

To quantify the performance of the SB-MKO model and, more generally, the impact of simplifying foot kinematics using a reduced number of synergies, we evaluated the translational and rotational errors, *e*_*t*_ and *e*_*r*_ , between experimental measurements and model estimates, considering foot and ankle separately. These errors were defined as:

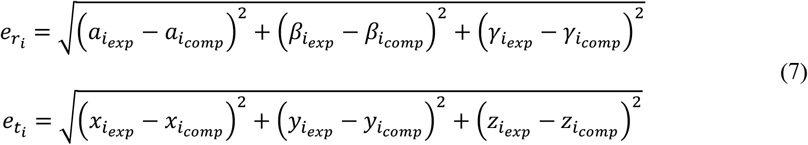

We first assessed the loss of information associated solely with reducing the number of synergies. For each specimen, foot postures were reconstructed using an increasing number of individual synergies, using the corresponding coefficients obtained from Eq. (6). This allowed us to evaluate how model accuracy degrades as fewer synergies are retained.

To characterize the overall accuracy of the complete SB-MKO process, a set of virtual markers was placed on each specimen in the neutral posture (Fig. 1). These markers were then propagated to all scanned postures using the rototranslational matrices of the corresponding bones. The generic synergy-based model was scaled to each specimen, and SB-MKO was applied to reconstruct all postures, allowing for the quantification of *e*_*t*_ and *e*_*r*_. To provide an additional, more direct comparison, the marker tracking error, i.e., the average absolute distance between experimental markers and those reconstructed via SB-MKO, was also computed in this case.

**Figure 1:**
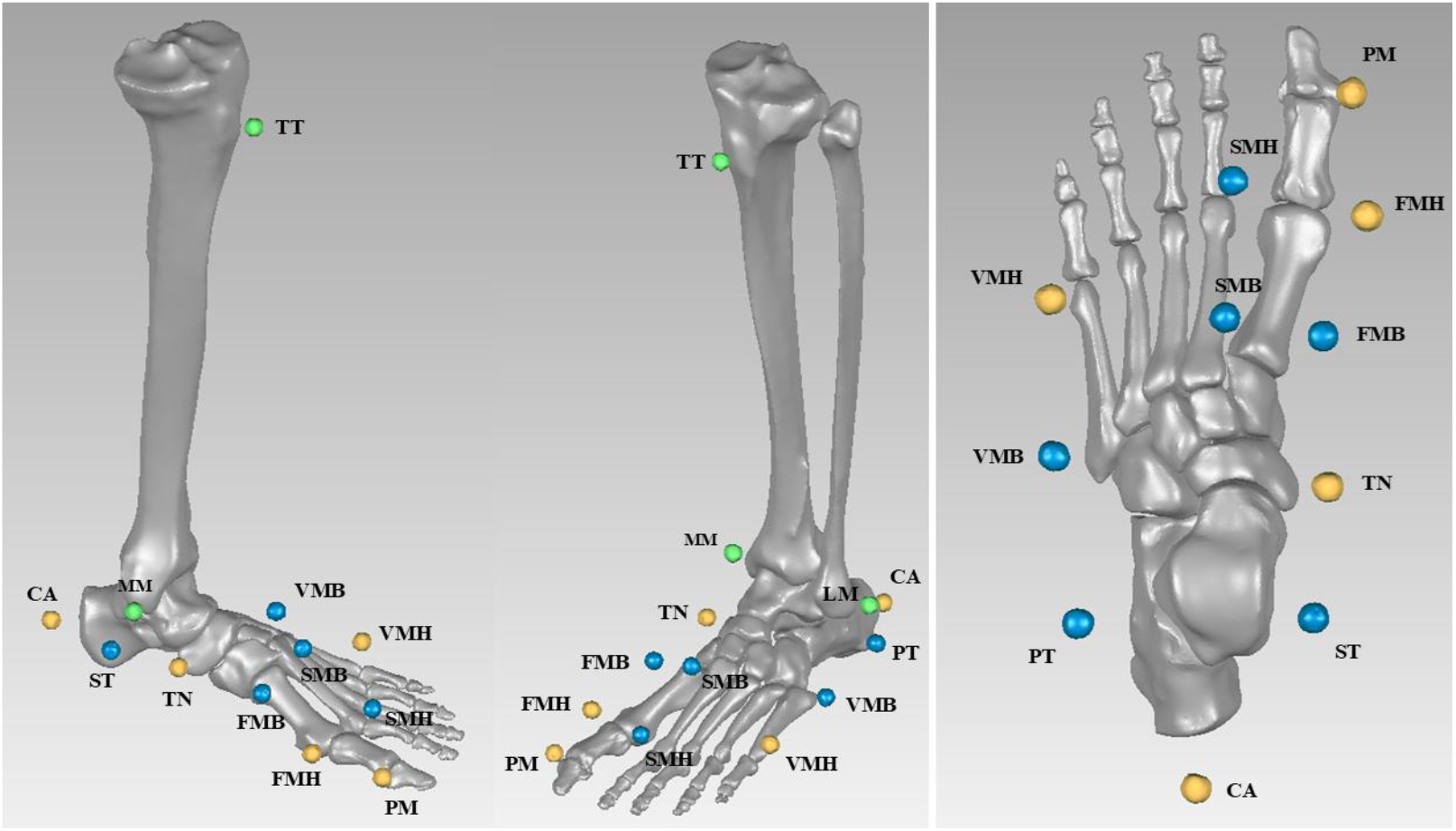
Virtual marker locations and naming. In green are depicted the markers belonging to the shank. On the right, a detailed view of the Rizzoli marker set [34]. In yellow, the markers belonging to the reduced set are highlighted.

Finally, to evaluate the robustness of SB-MKO with respect to a reduced number of foot markers, the same analysis was repeated using only four markers from the original Rizzoli protocol: CA, TN, FMH, and VMH.

### 2.5 Experimental test on clinical data

As a proof-of-concept of the new SB-MKO’s performance in a clinical context, we compared its ability to discriminate among different populations with a traditional MSFM (Leardini, et al., 2007). We considered three groups, healthy (3 individuals, 4 feet), flat (3 individuals, 6 feet) and diabetic (3 individuals, 6 feet). Population data are reported in Table 2. Each participant performed several gait trials at the Movement Analysis Lab at IOR (Leardini, et al., 2019).

**Table 2:** Average age and body mass index (BMI) for the three considered populations.

|  | Healthy | Flat | Diabetic |
| --- | --- | --- | --- |
| Number of feet | 4 | 6 | 6 |
| Age | 44.3 ± 23.2 | 50.9 ± 13.6 | 58.1 ± 12.0 |
| BMI | 22.6 ± 0.7 | 27.3 ± 4.0 | 29.4 ± 6.8 |

Once the foot kinematics was reconstructed with both approaches and using the full marker set (i.e., 10 markers), we computed three angles defining the foot arches: the medial longitudinal arch angle (MLA), defined as the angle between the x axis of the anatomical reference system of CA and M1 (Fig. 2.a); the lateral longitudinal arch angle (LLA), defined as the angle between the x axis of the anatomical reference system of CA and M5 (Fig. 2.b); the transverse arch angle (TRA), defined as the angle between the y axis of the anatomical reference system of CM and CU (Fig. 2.c). This latter was computed only within the SB-MKO, as MSFM treats the midfoot as a rigid segment, resulting in a constant value throughout the gait cycle.

**Figure 2:**
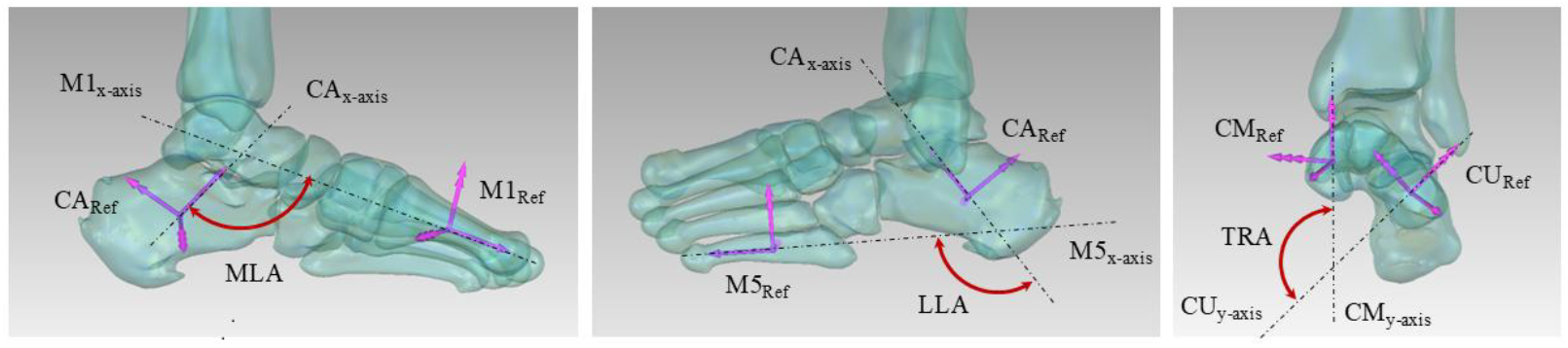
Graphical representation of the three angles considered to describe the foot arches: medio-longitudinal arch angle-MLA (left); lateral-longitudinal arch angle - LLA (center); transverse arch angle - TRA (right).

Finally, a t-test was performed (p<0.01) to identify differences between the groups, using the introduced measurement of foot arches.

## 3 RESULTS

### 3.1 Model accuracy

The quantification of the rotational and translational errors due to using a reduced number of individual synergies confirmed previous observations (Conconi, Sancisi, Leardini, & Belvedere, 2024). At the ankle, a single synergy results in an average rotational error of 1.14 ± 0.26° and in an average translational error of 0.46 ± 0.10 mm (Fig. 3). Conversely, at the foot three synergies resulted in an average rotational error of 0.88 ± 0.17° and in an average translational error 0.42 ± 0.05 mm (Fig. 3). Overall these results support the use of only four synergies to describe the motion of the foot and ankle complex, when toes are not considered.

**Figure 3:**
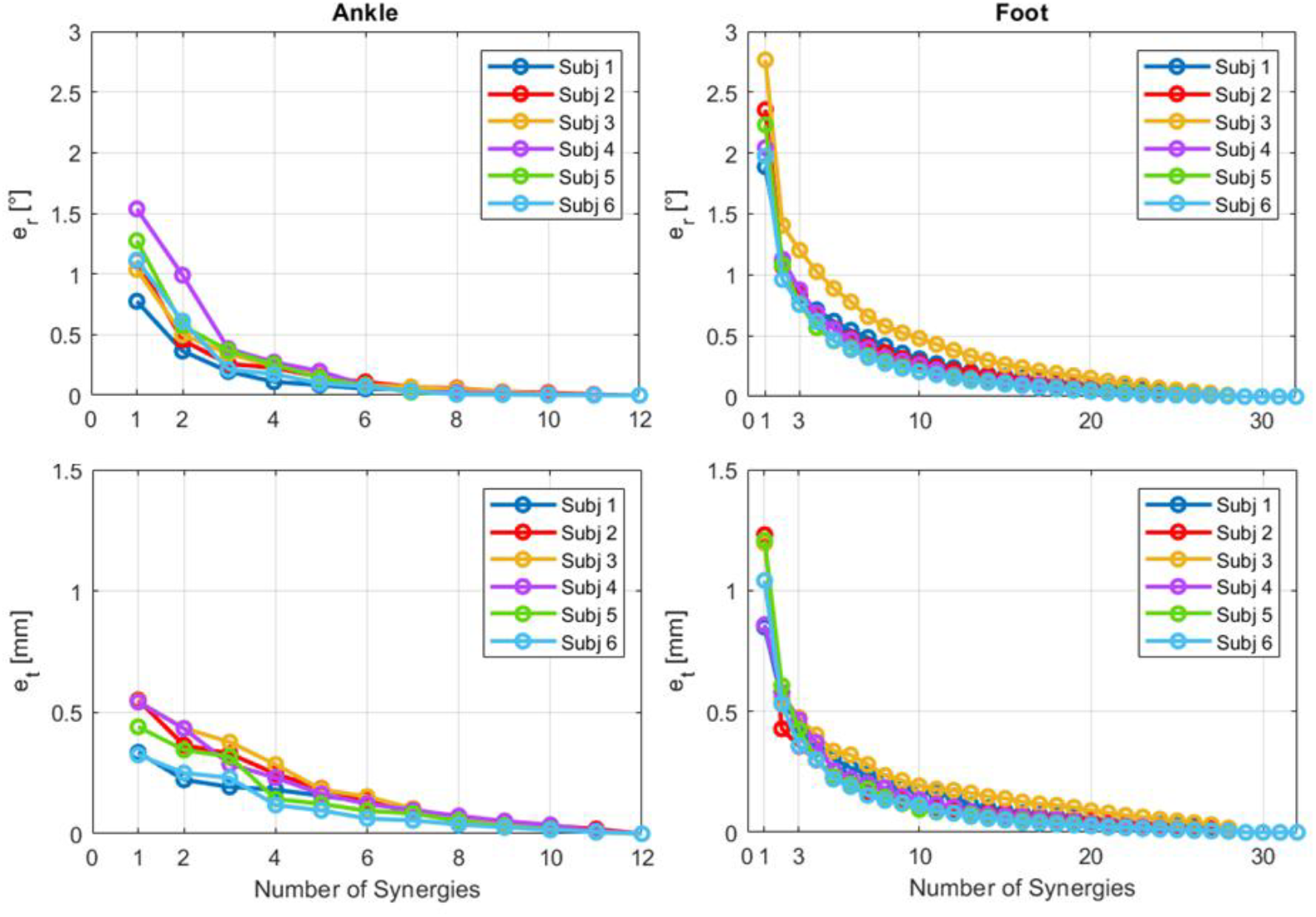
Rotational (e_r_, first row) and translational (e_t_, second row) errors at the ankle (left column) and at the foot (right column) as a function of the synergy number adopted for the posture reconstruction for the three considered subjects.

The SB-MKO accuracy was quantified in average errors of 4.52 ± 1.63 ° and 3.86 ± 2.28 mm, and of 6.83 ± 0.86 ° and 3.48 ± 0.48 mm for the ankle and the foot, respectively (Table 3, Fig. S1), with a marker residual error of 0.88 ± 0.44 mm (fig. S4). This level of accuracy is in line or better than previous literature (Benoit, et al., 2006) (Nester, et al., 2007) (Okita, Meyers, Challis, & Sharkey, 2009) (Shultz, Kedgley, & Jenkyn, 2011) (Kessler, et al., 2019) (Schallig, et al., 2021).

**Table 3:** Rotational (e_r_) and translational (e_t_) errors at the ankle and at the foot of the analyzed specimens using SB-MKO, with the full and reduced foot maker set.

|  | Ankle |  |  |  | Foot |  |  |  |
| --- | --- | --- | --- | --- | --- | --- | --- | --- |
| | $e_r [^\circ]$ | | $e_t$ [mm] | | $e_r [^\circ]$ | | $e_t$ [mm] | |
|  | Full<br>marker<br>set | Reduced<br>marker<br>set | Full<br>marker<br>set | Reduced<br>marker<br>set | Full<br>marker<br>set | Reduced<br>marker<br>set | Full<br>marker<br>set | Reduced<br>marker<br>set |
| Subject 1 | $2.83 \pm 0.56$ | $2.69 \pm 0.61$ | $2.30 \pm 0.15$ | $2.31 \pm 0.15$ | $6.78 \pm 0.47$ | $7.36 \pm 0.55$ | $3.66 \pm 0.31$ | $4.06 \pm 0.37$ |
| Subject 2 | $6.51 \pm 0.35$ | $5.93 \pm 0.28$ | $2.47 \pm 0.22$ | $2.46 \pm 0.23$ | $7.74 \pm 0.41$ | $7.48 \pm 0.47$ | $3.54 \pm 0.35$ | $3.36 \pm 0.29$ |

Finally, using a reduced marker set resulted in rotational and translational errors of 4.23 ± 1.40° and 3.86 ± 2.28 mm at the ankle, and 7.25 ± 0.78° and 3.62 ± 0.29 mm at the foot (Table 3, Fig. S4), with a marker residual error of 1.03 ± 0.51 mm (Fig. S5). Thus, the substantial experimental simplification enabled by SB-MKO does not significantly affect the accuracy of gait analysis results.

### 3.2 Experimental test on clinical data

MLA reconstructed by means of standard MSFM allows differentiation between healthy and flat feet only during the late phase of the push-off and early swing (fig. 4). No differences were observable between healthy and diabetic feet, while the latter differed from flat feet over the entire step. Very similar conclusions can be drawn regarding the LLA.

**Figure 4:**
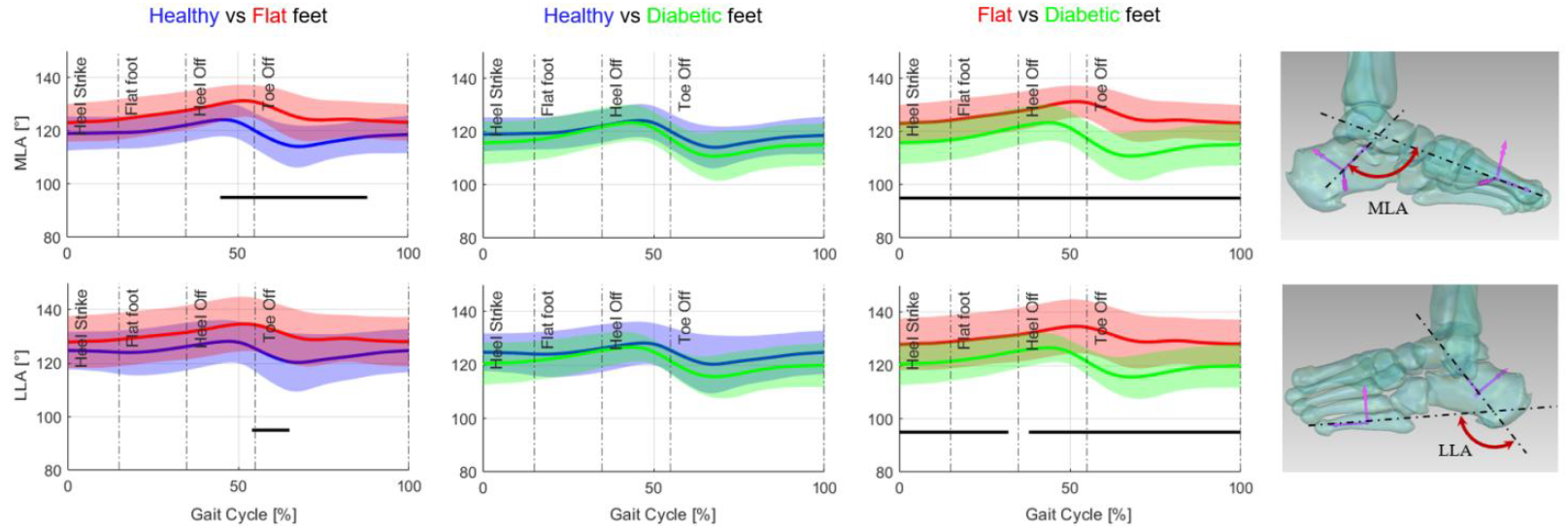
Comparison of the medial longitudinal arch angle - MLA (top row) and the lateral longitudinal arch angle - LLA (bottom row) between healthy, diabetic, and flat feet individuals using the standard MSFM approach. Black lines denote statistically significant differences between populations.

In contrast, when observing the same angles using the SB-MKO approach, the differences between the groups are much more evident (Fig. 5). Healthy and flat feet differ statistically across the entire step cycle for the MLA, while the other comparisons are significant over more than 85% of the cycle. Contrary to what observed through MSFM, LLA revealed in this case differences between healthy and flat feet and not between flat and diabetic feet. The TRA, whose variation cannot be quantified within the rigid cluster typical of MSFM, showed a power of discrimination similar to MLA.

**Figure 5:**
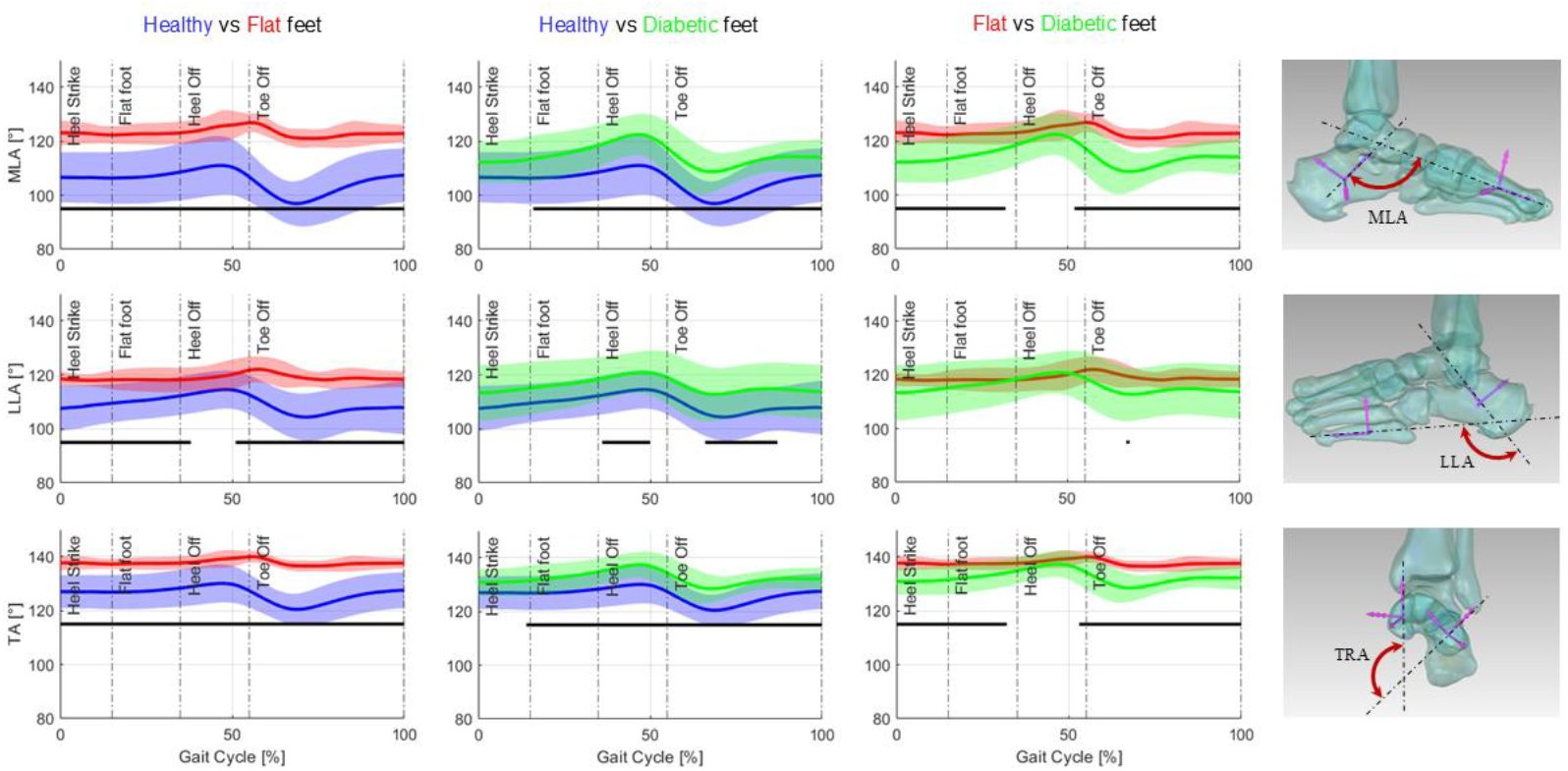
Comparison of the medial longitudinal arch angle – MLA (top row), lateral longitudinal arch angle - LLA (middle row), and transverse arch angle - TRA (bottom row) between healthy, diabetic, and flat feet individuals using the SB-MKO approach. Black lines denote statistically significant differences between populations.

In general, the SB-MKO resulted in a smaller standard deviation with respect to the MSFM, confirming its capability to filter the soft tissue and maker occlusion artefacts.

## 4 DISCUSSION

In this work, we presented a new synergy-based multibody model of the foot and ankle, capable of reconstructing the three-dimensional kinematics of all bones in the articular complex using a five-DOF system. This is achieved by exploiting intrinsic couplings among bone motion components, i.e., motion synergies, experimentally identified through principal component analysis.

The present analysis confirmed that the representation of foot and ankle motion, excluding the toes, with four DOFs can be accurate, yielding rotational and translational errors of about 1° and 0.5 mm when using subject-specific synergies. Similarly, when the generic SB-MKO model is used to reconstruct the experimental postures, the average rotational and translational errors resulted in 4.52 ± 1.63° and 3.86 ± 2.28 mm for the ankle and 6.83 ± 0.86 ° and 3.48 ± 0.48 mm for the foot, due to the additional effect of averaging and scaling the model. The corresponding tracking errors between model and experimental markers were always below 1.00 mm. This level of accuracy is in line with or better than previous literature (Benoit, et al., 2006) (Nester, et al., 2007) (Okita, Meyers, Challis, & Sharkey, 2009) (Shultz, Kedgley, & Jenkyn, 2011) (Kessler, et al., 2019) (Schallig, et al., 2021), thus supporting the applicability of the proposed approach for the analysis of foot biomechanics.

From a clinical perspective, the SB-MKO model demonstrated greater ability to discriminate among groups than the standard MSFM approach when applied to gait data. The MSFM was able to distinguish healthy from pathological feet only during late push-off and early swing, and did not detect differences between healthy and diabetic individuals (Fig. 4). In contrast, the SB-MKO clearly differentiated all three populations and revealed higher arch mobility in healthy subjects compared with individuals with flat feet (Fig. 5). Within-group variability was also reduced, reflecting the filtering effect of the rigid kinematic chain on soft-tissue and marker occlusion artefacts. Furthermore, access to intersegmental mobility made it possible, for the first time to the authors’ knowledge, to examine transverse arch deformation, which again showed significant differences among groups. These results highlight the importance of capturing bone-level kinematics, which cannot be achieved with standard segmental models.

The limited total number of DOF, five for the ankle, foot, and toes motion, enables a substantial reduction in the number of markers required to reconstruct foot kinematics during gait analysis. Our results showed that only four of the ten markers in the Rizzoli protocol are sufficient to describe the foot kinematics, with no meaningful loss of accuracy. Using the reduced marker set, rotational and translational errors were 4.23 ± 1.40° and 3.86 ± 2.28 mm at the ankle, and 7.25 ± 0.78 ° and 3.62 ±0.29 mm at the foot, showing negligible differences compared with the full marker set. This reduction offers several advantages for gait analysis. The experimental protocol becomes simpler and faster to implement; fewer markers reduce the likelihood of marker overlapping on camera sensors, thereby decreasing the impact of MOA; and, aided by MKO filtering, the reconstructed kinematics become less sensitive to the low repeatability of foot marker placement reported in the literature (Di Marco, Rossi, Racic, Cappa, & Mazzà, 2016). Finally, the robustness of the SB-MKO to the number of foot markers makes the method easily applicable across different protocols. Existing datasets can be retrospectively analyzed and compared, even when different marker sets were employed, thereby improving data sharing by harmonizing historical gait analysis data (Besier, et al., 2024). On this point, it is worth noting that the reduced marker set considered here is a subset of the one used in the Rizzoli protocol, but other reduced marker sets may improve accuracy and robustness.

The present analysis has limitations. Synergies were derived from a small number of non-pathological specimens. This limitation is inherent to the experimental complexity of measuring spatial kinematics for all foot bones, while the radiation exposure associated with extensive scanning precludes in-vivo experiments. Despite the small sample size, the high correlation among synergies indicates that foot mobility is largely consistent across individuals. The model accuracy has been determined on static scans: this provides an adequate gold standard for quantifying pose error due to synergy reduction, averaging, and scaling, while soft-tissue and marker occlusion artifacts did not affect the comparison results. Accuracy may therefore decrease when applied to standard gait data. The model was applied to both healthy and pathological populations, even though synergies may change with disease onset or progression, or in the presence of bone deformities, soft tissue deficiencies, or alterations. Nevertheless, the model successfully discriminated among groups, suggesting that the simplification is acceptable at least for the conditions examined. The synergy describing toe motion was derived from a geometric approximation of the metatarsophalangeal joints rather than experimental data, due to limits imposed by scanner dimension, and inter-phalangeal motion was not modeled. Toe kinematics may therefore be captured with reduced accuracy. Finally, the adopted synergies have been derived under a weight-bearing scenario that represents a limited subset of the loads experienced by the foot and ankle during functional tasks of daily living, both externally, due to environmental loads, and internally, due to muscle activations. It is therefore possible that more synergies would be needed to fully describe the foot behavior during dynamic tasks such as running or jumping.

Beyond kinematic reconstruction, the proposed model provides a solid basis for a more accurate investigation of foot–ground interaction. By capturing intersegmental mobility within a mechanically consistent rigid multibody formulation, the model enables more reliable identification of which foot bones are in contact with the ground at a given instant and how contact conditions evolve during motion. This, in turn, allows ground reaction forces to be redistributed among individual foot bones in a more physiologically meaningful manner than is possible with rigid or weakly articulated foot representations. The resulting bone-level contact force estimates can serve as anatomically consistent inputs for subsequent deformable or finite-element models of the foot, where they may be applied as boundary conditions to investigate internal stresses and tissue loading. Moreover, the limited number of generalized coordinates and the absence of redundant degrees of freedom make the formulation particularly well suited for forward dynamic and predictive simulations, in which numerical conditioning, stability, and computational efficiency are critical, together with a robust and interpretable representation of foot–ground contact.

The current model is publicly available in both a MATLAB implementation, which includes a C++ based visualizer (Conconi, 2026), and an OpenSim implementation (Modenese, 2026), both currently limited to kinematic analysis. Future work will extend the model by integrating it into a full lower-limb musculoskeletal framework, incorporating intrinsic foot muscles and ligaments, and adding a contact model to evaluate how ground reaction forces are distributed across the bones. The experimental determination of foot synergies will be extended to additional specimens, possibly investigating the effects of foot pathologies and, eventually, the foot kinematics in vivo through dynamic MRI (Conconi, et al., 2023).

## AUTHORS’ CONTRIBUTIONS

MC participated in the design of the study, the derivation of the model, its numerical implementation, and contributed to data collection and analysis as well as writing of the manuscript; LM contributed in the numerical implementation of the model, in data analysis, as well as writing of the manuscript; GMB contributed the numerical investigation of model accuracy and robustness and in the data analysis; AL and CB contributed to design the experimental setup, to the data collection and analysis, as well as in the writing of the manuscript; NS contributed to the design of the study and the derivation of the model, as well as in the writing of the manuscript. All authors have read and approved the final version of the manuscript, and agree with the order of presentation of the authors.

## SUPPLEMENTARY MATERIALS

Supplementary materials associated with this article can be found in the online version.

