## Supplementary material for "A reduced-order multibody model of the foot–ankle complex based on kinematic synergies"

### A novel, synergy-based kinematic model of the foot and ankle for gait analysis: supplementary material

**Parametrization of the bone pose and foot posture**

Bone 3D orientation was expressed through a variation of the classic Grood & Suntay cardanic sequence [45] optimized for the foot: considering the relative motion of a distal with respect to a proximal bone, pronation/supination (α) is the rotation about the x axis of the distal bone, dorsi/plantar flexion (γ) is the rotation about the z axis of the proximal bone, abduction/adduction (β) is the rotation about the floating axis, i.e. perpendicular to the previous two. Bone position was represented by means of the three coordinates of the origin **o** of the moving ARS. Hereinafter, bold letters denote vectors, bold capital letters denote matrixes, c and s represent cosine and sine functions, respectively. The roto-translation matrix representing the pose of a bone will therefore be

$$\boldsymbol{M} = \boldsymbol{T}(\bar{O})\boldsymbol{R}_z(\gamma)\boldsymbol{R}_y(\beta)\boldsymbol{R}_x(a) =$$

$$= \begin{bmatrix} \mathrm{c}\,(\gamma)\mathrm{c}(\beta) & (\mathrm{c}(\gamma)\,\mathrm{s}(\beta)\mathrm{s}(a) - \mathrm{s}(\gamma)\,\mathrm{c}(a)) & (\mathrm{c}(\gamma)\,\mathrm{s}(\beta)\mathrm{c}(a) + \mathrm{s}(\gamma)\,\mathrm{s}(a)) & \mathrm{o}_x \\ \mathrm{s}\,(\gamma)\mathrm{c}(\beta) & (\mathrm{s}(\gamma)\,\mathrm{s}(\beta)\mathrm{s}(a) + \mathrm{c}(\gamma)\,\mathrm{c}(a)) & (\mathrm{s}(\gamma)\,\mathrm{s}(\beta)\mathrm{c}(a) - \mathrm{c}(\gamma)\,\mathrm{s}(a)) & \mathrm{o}_y \\ -\,\mathrm{s}(\beta) & \mathrm{c}(\beta)\mathrm{s}(a) & \mathrm{c}(\beta)\mathrm{c}(a) & \mathrm{o}_z \\ 0 & 0 & 0 & 1 \end{bmatrix}$$

where $\boldsymbol{R}_i$ is the matrix corresponding to a pure rotation about the i-th axis and $\boldsymbol{T}$ to a pure translation. Conversely, rotational and translational parameters can be extracted from each matrix as

$$a = tan^{-1}(\boldsymbol{M}(3{,}2)/\boldsymbol{M}(3{,}3))$$
$$\beta = tan^{-1}\left(\boldsymbol{M}(3{,}1)/\sqrt{\boldsymbol{M}(3{,}2)^2 + \boldsymbol{M}(3{,}3)^2}\right)$$
$$\gamma = tan^{-1}(\boldsymbol{M}(2{,}1)/\boldsymbol{M}(1{,}1))$$
$$x = \boldsymbol{M}(1{,}4)$$
$$y = \boldsymbol{M}(1{,}5)$$
$$z = \boldsymbol{M}(1{,}6)$$

where the domain of the arctangent can be extended to 2π considering the sign of the elements in the matrix. The pose vector associated with the matrix $\boldsymbol{M}$ can be expressed as

$$\boldsymbol{p}_{\mathrm{i}} = [a_i, \beta_{\mathrm{i}}, \gamma_{\mathrm{i}}, x_{\mathrm{i}}, y_{\mathrm{i}}, x_{\mathrm{i}}]$$

This parametrization holds independently from the reference system in which the bone pose is expressed. In particular, in this work we express the position of all the bone in the foot and ankle complex relatively to another bone in the grup but the tibia, for which the pose with respect to the ground is considered. Specifically, the motion of talus (TA) and fibula (FI) were expressed relative to the tibia (TI) absolute pose; the motion of the calcaneus (CA) and the navicular (NA) relative to TA; the motion of the remaining bones (CU, cuboid; CM, medial cuneiform; CI, intermediate cuneiform; CL, lateral cuneiform; M1–5, first to fifth metatarsals) relative to NA.

In this context, to provide an example the relative pose of the talus in the tibia is represented by

$$\boldsymbol{M}_{TATI_i} = \boldsymbol{M}_{TI_i}^{-1}\boldsymbol{M}_{TA_i},\ i = 1, \cdots, n$$

where each matrix on the right side is the bone pose in the ground reference system, and $n = 32$ is the number of available foot posture. From here, the foot posture vector **p**, 108x1 vector, contains the concatenated positional and orientational coordinates of all the bones, so that

$$\boldsymbol{p} = [\boldsymbol{p}_{TATI}, \boldsymbol{p}_{FITI}, \boldsymbol{p}_{CATA}, \boldsymbol{p}_{NATA}, \boldsymbol{p}_{CUNA}, \boldsymbol{p}_{CMNA}, \boldsymbol{p}_{CINA}, \boldsymbol{p}_{CLNA}, \boldsymbol{p}_{M1NA}, \cdots, \boldsymbol{p}_{M5NA}]'$$

#### Synergies analysis

The analysis of cumulated variance for the ankle and foot synergies (Figure S1), support the model of the foot as a 4 DOF system, 1 at the ankle and three belonging to the foot itself.

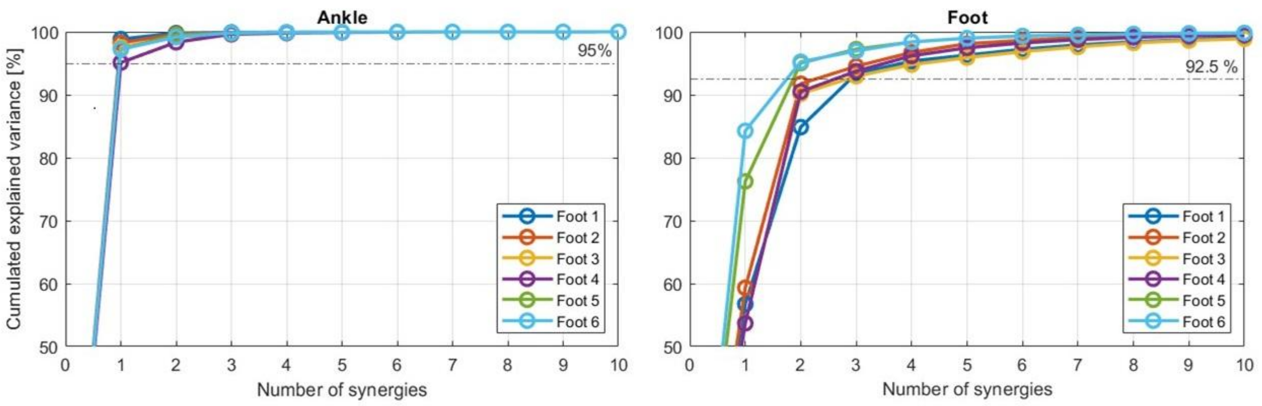


*Figure S1: Cumulated variance with the increasing number of considered synergies at the ankle (left) ant the foot (right)*

Synergies can be represented by plotting the amplitude of their individual components. Components, in which each sextuple will represent the angular and translational motion coupling of a specific bone. In figure S2 are reported the graphical representation of the 4 foot and ankle synergies for all the feet of this study.

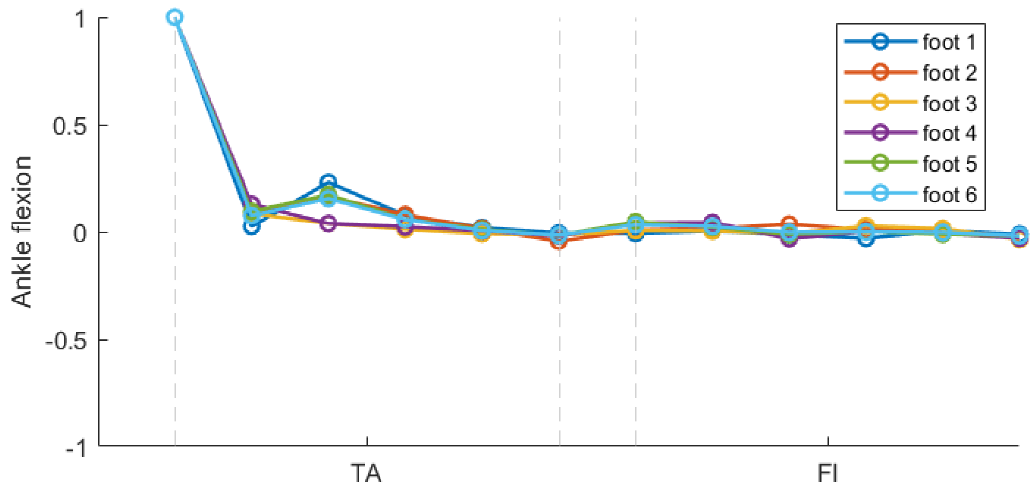


*Figure S2: Graphical comparison among the first ankle synergy for the six considered subjects*

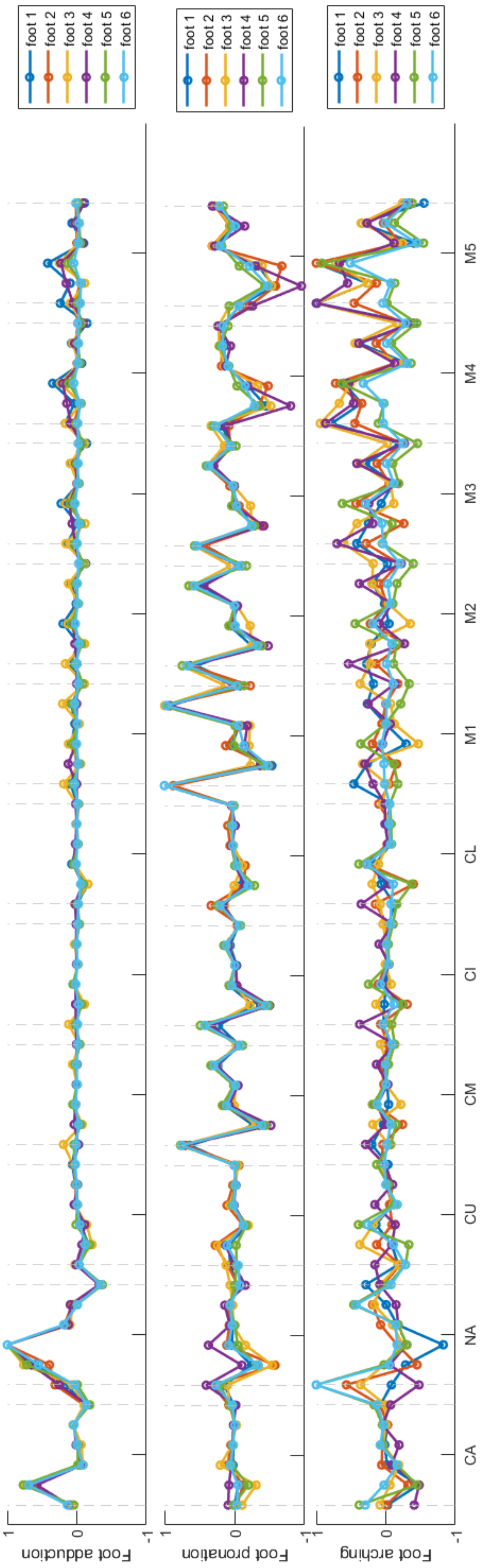


*Figure s3: Graphical comparison among the first three foot synergies for the six considered subjects*

**Reconstruction of foot posture through synergies**

The toes are here modeled as a set of five additional bodies, each coupled with the corresponding metatarsal through a hinge, whose rotation nis rigidly coupled with the big toe flexion.

In order to reconstruct the generic foot posture through synergies, we resort to the average posture and the average foot and ankle synergies

$$\boldsymbol{u} = [\boldsymbol{u}_{\mathrm{TATI}}, \boldsymbol{u}_{\mathrm{FITI}}, \boldsymbol{u}_{\mathrm{CATA}}, \boldsymbol{u}_{\mathrm{NATA}}, \boldsymbol{u}_{\mathrm{CUNA}}, \boldsymbol{u}_{\mathrm{CMNA}}, \boldsymbol{u}_{\mathrm{CINA}}, \boldsymbol{u}_{\mathrm{CLNA}}, \boldsymbol{u}_{\mathrm{M1NA}}, \cdots, \boldsymbol{u}_{\mathrm{M5NA}}, \boldsymbol{u}_{\mathrm{T1M1}}, \cdots, \boldsymbol{u}_{\mathrm{T5M5}}]$$

$$\boldsymbol{s}_1 = \left[\boldsymbol{s}_{\mathrm{TATI}_1}, \boldsymbol{s}_{\mathrm{FITI}_1}, \boldsymbol{0}_6, \boldsymbol{0}_6, \boldsymbol{0}_6, \boldsymbol{0}_6, \boldsymbol{0}_6, \boldsymbol{0}_6, \boldsymbol{0}_6, \cdots, \boldsymbol{0}_6, \boldsymbol{0}_6, \cdots, \boldsymbol{0}_6\right]$$

$$\boldsymbol{s}_i = \left[\boldsymbol{0}_6, \boldsymbol{0}_6, \boldsymbol{s}_{\mathrm{CATA}_i}, \boldsymbol{s}_{\mathrm{NATA}_i}, \boldsymbol{s}_{\mathrm{CUNA}_i}, \boldsymbol{s}_{\mathrm{CMNA}_i}, \boldsymbol{s}_{\mathrm{CINA}_i}, \boldsymbol{s}_{\mathrm{CLNA}_i}, \boldsymbol{s}_{\mathrm{M1}NA_i}, \cdots, \boldsymbol{s}_{\mathrm{M5NA}_i}, \boldsymbol{0}_6, \cdots, \boldsymbol{0}_6\right] \quad , with\ i = 2,3,4$$

$$\boldsymbol{s}_5 = \left[\boldsymbol{0}_6, \boldsymbol{0}_6, \boldsymbol{0}_6, \boldsymbol{0}_6, \boldsymbol{0}_6, \boldsymbol{0}_6, \boldsymbol{0}_6, \boldsymbol{0}_6, \boldsymbol{0}_6, \cdots, \boldsymbol{0}_6, \boldsymbol{s}_{\mathrm{T1}M1_5}, \cdots, \boldsymbol{s}_{\mathrm{T5M5}_5}\right]$$

where $\boldsymbol{0}_6 = [0,0,0,0,0,0]$.

It is possible to notice that the average posture present no motion of the TI, and that the synergies are decoupled, the first describing the ankle motion, the second to fourth the foot motion, and the fifth the toes motion.

The foot posture can thus be parametrized using 5 variables, namely the five synergy coefficients $\sigma_i$, as

$$\boldsymbol{p}(\boldsymbol{x}) = \boldsymbol{u} + \sum_{i=1}^{5} \sigma_i \boldsymbol{s}_i,$$

If $\boldsymbol{p}_{\mathrm{TI}}$ is the tibia pose with respect to the global observer, the matrixes describing the pose of each bone with respect to the same global observer can be obtained computing the product of relative pose matrixes, i.e.

$$\boldsymbol{M}_{TI} = \boldsymbol{M}(\boldsymbol{p}_{\mathrm{TI}})$$

$$\boldsymbol{M}_{TA} = \boldsymbol{M}(\boldsymbol{p}_{\mathrm{TI}})\, \boldsymbol{M}(\boldsymbol{p}_{\mathrm{TATI}})$$

$$\boldsymbol{M}_{FI} = \boldsymbol{M}(\boldsymbol{p}_{\mathrm{TI}})\, \boldsymbol{M}(\boldsymbol{p}_{\mathrm{FITI}})$$

$$\boldsymbol{M}_{CA} = \boldsymbol{M}(\boldsymbol{p}_{\mathrm{TI}})\, \boldsymbol{M}(\boldsymbol{p}_{\mathrm{TATI}})\boldsymbol{M}(\boldsymbol{p}_{\mathrm{CATA}})$$

$$\boldsymbol{M}_{NA} = \boldsymbol{M}(\boldsymbol{p}_{\mathrm{TI}})\, \boldsymbol{M}(\boldsymbol{p}_{\mathrm{TATI}})\, \boldsymbol{M}(\boldsymbol{p}_{\mathrm{NATA}})$$

$$\boldsymbol{M}_{CU} = \boldsymbol{M}(\boldsymbol{p}_{\mathrm{TI}})\, \boldsymbol{M}(\boldsymbol{p}_{\mathrm{TATI}})\, \boldsymbol{M}(\boldsymbol{p}_{\mathrm{NATA}})\boldsymbol{M}(\boldsymbol{p}_{CUNA})$$

$$\boldsymbol{M}_{CM} = \boldsymbol{M}(\boldsymbol{p}_{\mathrm{TI}})\, \boldsymbol{M}(\boldsymbol{p}_{\mathrm{TATI}})\, \boldsymbol{M}(\boldsymbol{p}_{\mathrm{NATA}})\, \boldsymbol{M}(\boldsymbol{p}_{CMNA})$$

$$\boldsymbol{M}_{CI} = \boldsymbol{M}(\boldsymbol{p}_{\mathrm{TI}})\, \boldsymbol{M}(\boldsymbol{p}_{\mathrm{TATI}})\, \boldsymbol{M}(\boldsymbol{p}_{\mathrm{NATA}})\, \boldsymbol{M}(\boldsymbol{p}_{CINA})$$

$$\boldsymbol{M}_{CL} = \boldsymbol{M}(\boldsymbol{p}_{\mathrm{TI}})\,\boldsymbol{M}(\boldsymbol{p}_{\mathrm{TATI}})\,\boldsymbol{M}(\boldsymbol{p}_{\mathrm{NATA}})\,\boldsymbol{M}(\boldsymbol{p}_{CLNA})$$

$$\boldsymbol{M}_{Mi} = \boldsymbol{M}(\boldsymbol{p}_{\mathrm{TI}})\,\boldsymbol{M}(\boldsymbol{p}_{\mathrm{TATI}})\,\boldsymbol{M}(\boldsymbol{p}_{\mathrm{NATA}})\,\boldsymbol{M}(\boldsymbol{p}_{MiNA}), with\ i = 1, \cdots, 5$$

$$\boldsymbol{M}_{Ti} = \boldsymbol{M}(\boldsymbol{p}_{\mathrm{TI}})\,\boldsymbol{M}(\boldsymbol{p}_{\mathrm{TATI}})\,\boldsymbol{M}(\boldsymbol{p}_{\mathrm{NATA}})\,\boldsymbol{M}(\boldsymbol{p}_{MiNA})\boldsymbol{M}(\boldsymbol{p}_{TiMi}), with\ i = 1, \cdots, 5$$

**Anisotropic scaling**

The foot and ankle complex can differ in width, length and height from subject to subject, so anisotropic scaling may provide a better fit of a generic model to the specificity of a given individual respect to isotropic one.

Due to our anatomical definition of the reference systems, their axes are in general not parallel to each other, making anisotropic scaling not straightforward. Indeed, anisotropic scaling do not preserve orthogonality on directions different from the principal ones. For this reason we adopt an approximated approach consisting in several steps.

We assume, as experimentally observed, that the average foot posture can be considered as a good approximation of the neutral one, at least for scaling purposes. As described in the previous section, we computed the global rototranslational matrixes of each bone at the average posture and use the same to determine the location of all the virtual marker, whose coordinates are originally expressed in the ARS of the bone to which they are attached.

With this foot posture and virtual marker distribution, we computed the scaling factor for each axis by comparing maker distance in the static pose for the considered subject and in the average pose of the generic foot model. The markers name are defined according to [1] and are so distributed on the bones:

- MM, LM, and TT in TI;
- CA, PT, and ST in CA;
- TN in NA;
- FMB and FMH in M1;
- SMB and SMH in M2;
- VMB and VMH in M5;
- PM in T1

The detailed procedure is available as a MATLAB script, commented in the details, at [2].

*Tibia scaling factors*

The tibia is associated with three markers, laying on a coronal plane. To determine the scaling factor in the proximo/distal direction (y axis of TI ARS), we evaluate the average distance between the marker on each malleolus and the one on the tibial tuberosity. The scaling in the medio/lateral direction (z axis of TI ARS) was defined based on the inter-malleolar distance. Finally the antero/posterior scaling (x axis of the TI ARS) was calculated as the average of the previous two. Mathematically, denoting with an *s* maker from the static scan and with an *a* marker from the average posture:

$$k_{y_{TI}} = \frac{(\|TT_s - MM_s\| + \|TT_s - ML_s\|)/2}{(\|TT_a - MM_a\| + \|TT_a - ML_a\|)/2}$$

$$k_{z_{TI}} = \frac{\|LM_s - MM_s\|}{\|LM_a - MM_a\|}$$

$$k_{x_{TI}} = \left(k_{y_{TI}} + k_{z_{TI}}\right)/2$$

$$\boldsymbol{S}_{TI} = \begin{bmatrix} k_{x_{TI}} & 0 & 0 \\ 0 & k_{y_{TI}} & 0 \\ 0 & 0 & k_{z_{TI}} \end{bmatrix}$$

*Foot scaling factors*

The bones in the foot are scaled in antero/posterior direction (x axis of the average foot posture) using the average distance between the calcaneal posterior marker and metatarsal head markers on M1 and M5. This three marker was also used to define an approximated foot contact plane orientation and its normal *n*. The scaling in the proximo/distal direction (y axis of the average foot posture) was obtained averaging the projection on the foot plane normal *n* of the vectors from the calacaneal posterior marker to the two malleolar ones. Finally the medio/lateral scaling was obtained looking ad the distance between the first and fifth metatarsal head markers. Mathematically:

$$k_{x_{foot}} = \frac{(\|FMH_s - CA_s\| + \|VMH_s - CA_s\|)/2}{(\|FMH_a - CA_a\| + \|VMH_a - CA_a\|)/2}$$

$$k_{y_{foot}} = \frac{\big(\boldsymbol{n} \cdot (LM_s - CA_s) + \boldsymbol{n} \cdot (MM_s - CA_s)\big)/2}{\big(\boldsymbol{n} \cdot (LM_a - CA_a) + \boldsymbol{n} \cdot (MM_a - CA_a)\big)/2}$$

$$k_{z_{foot}} = \frac{\|FMH_s - VMH_s\|}{\|FMH_a - VMH_a\|}$$

$$\boldsymbol{S}_{foot} = \begin{bmatrix} k_{x_{foot}} & 0 & 0 \\ 0 & k_{y_{foot}} & 0 \\ 0 & 0 & k_{z_{foot}} \end{bmatrix}$$

*Toes scaling factor*

Due to the lack of markers on the toes but the one on the big one, isotropic scaling was considered. In particular, toes scaling was guided by the distance between the markers on the big toe and the first metatarsal head, namely:

$$k_{toes} = \frac{\|PM_s - FMH_s\|}{\|PM_a - FMH_a\|}$$

$$\boldsymbol{S}_{toes} = \begin{bmatrix} k_{toes} & 0 & 0 \\ 0 & k_{toes} & 0 \\ 0 & 0 & k_{toes} \end{bmatrix}$$

*Scaling procedure*

Markers centers, bone stl models, and origin of ARS (all expressed in the average foot posture) are anisotropically scaled by multiplying them by the corresponding scaling matrix, for instance, given a set of points in the foot $\boldsymbol{P}_{foot}$ we will have:

$$\boldsymbol{P}_{foot,scaled} = \boldsymbol{S}_{foot} \cdot \boldsymbol{P}_{foot}$$

Since the orientation of the bone ARS is not parallel to the tibial one, anisotropic scaling would result in a non-orthogonal reference systems, corrupting the parametrization of the bone pose. For this reason, ARS orientation was preserved, assuming its variation to be negligible in this context.

Subsequently, synergies and are scaled and for the same reason only the translational components are considered. Moreover, as these latter represent translation about axes that in general are not parallel to the ones in which anisotropic scaling was computed, an isotropical approach was employed. As synergies describe relative articular motion,  scaling factor are determined as the average among those of the articulating bones. Mathematically, the synergy scaling factors are:

$$k_{s_1} = \left(k_{x_{TI}} + k_{y_{TI}} + k_{z_{TI}} + k_{x_{foot}} + k_{y_{foot}} + k_{z_{foot}}\right)/6$$

$$k_{s_i} = \frac{\left(k_{x_{foot}} + k_{y_{foot}} + k_{z_{foot}}\right)}{3}, for\ i = 2,3,4$$

$$k_{s_5} = \left(k_{x_{foot}} + k_{y_{foot}} + k_{z_{foot}} + 3 * k_{toes}\right)/6$$

After scaling, maker centers are back projected in the zero pose of the corresponding bones by computing the rototranslational matrixes as described in the previous section, using however the scaled average posture, and inverting them. Mathematically, the new scaled coordinates of the generic marker $\boldsymbol{mk}_{scaled}$ will be

$$\begin{bmatrix} \boldsymbol{mk}_{scaled} \\ 1 \end{bmatrix} = \boldsymbol{M}(\boldsymbol{u}_{\boldsymbol{scaled}})^{-1} \begin{bmatrix} \boldsymbol{S} & \boldsymbol{0} \\ \boldsymbol{0} & 1 \end{bmatrix} \boldsymbol{M}(\boldsymbol{u}) \begin{bmatrix} \boldsymbol{mk} \\ 1 \end{bmatrix}$$

The so scaled model is then fitted on the static trial via SB-MKO: an optimization is run that compute the vector $\boldsymbol{x}$**,** which parametrizes the foot and ankle posture with the 11 variables including the six components of $\boldsymbol{p}_{\mathrm{TI}}$ and the five synergy coefficients $\alpha_i$ , that minimize the sum of the squared distances between real and virtual markers. For the generic marker, we will thus compute

$$\left\| \begin{bmatrix} \boldsymbol{mk}_{exp} \\ 1 \end{bmatrix} - \boldsymbol{M}(\boldsymbol{p}(\boldsymbol{x})) \begin{bmatrix} \boldsymbol{mk} \\ 1 \end{bmatrix} \right\|^2$$

To compensate for variation in hallux abduction and lack of markers to capture the actual flexion of the various toes, the average posture of this latter is modified at this stage. In particular, the angle between the TI antero/posterior direction (x axis of TI ARS) and the line through the hallux and first metatarsal head experimental marker is computed  and the average posture is modified to match this angle. Likewise, the flexion values of the average posture of all the toes are modified so that the antero/posterior direction (x axis of Ti ARS) are laying the same plane of the one of the hallux.

Finally, to compensate for possible maker malposition, the experimental markers are back projected in the zero pose of the corresponding bone and substitute with the model makers for subsequent SB-MKO of dynamic trials.

**SB-MKO accuracy on the individual subjects and foot posture**

In what follow, the rotational and translational errors, $e_r$ and $e_t$ respectively, for the ankle and the foot at different postures are reported for each considered subject.

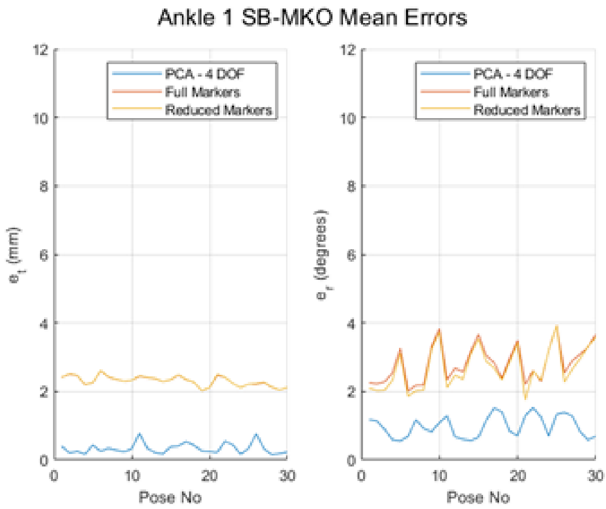


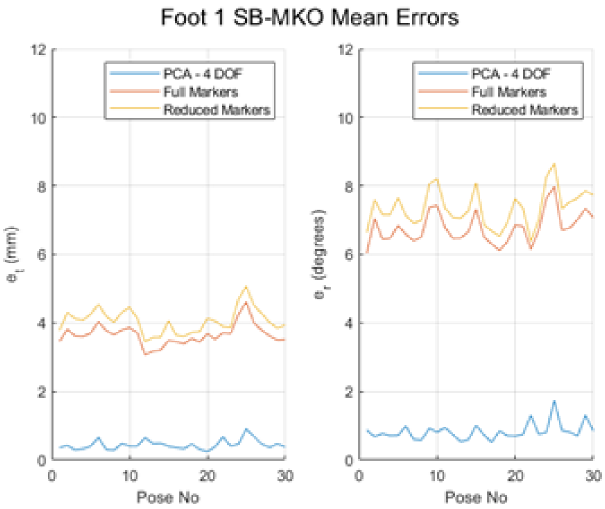


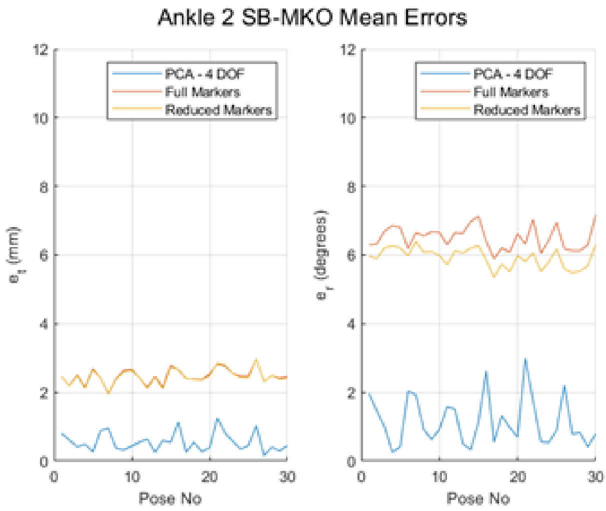


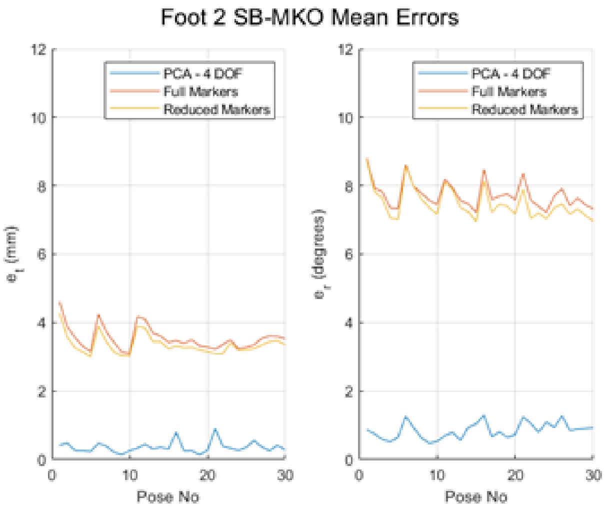


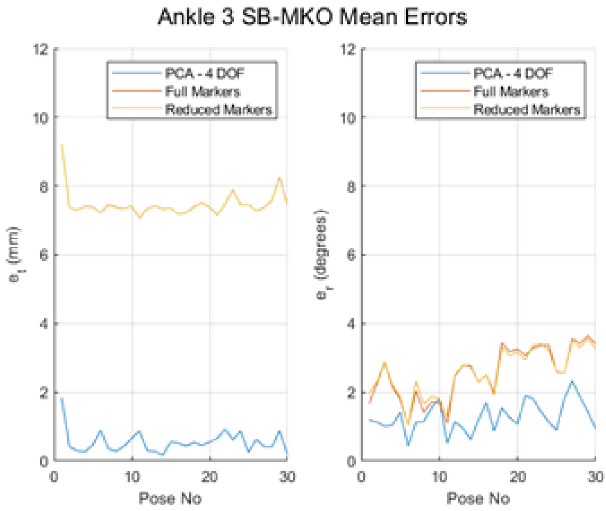


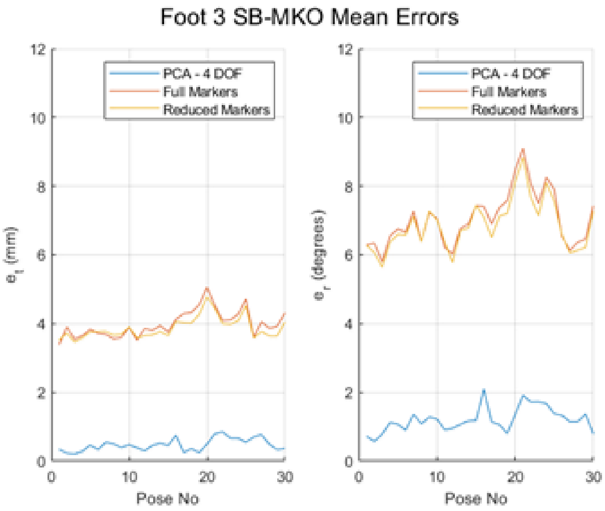

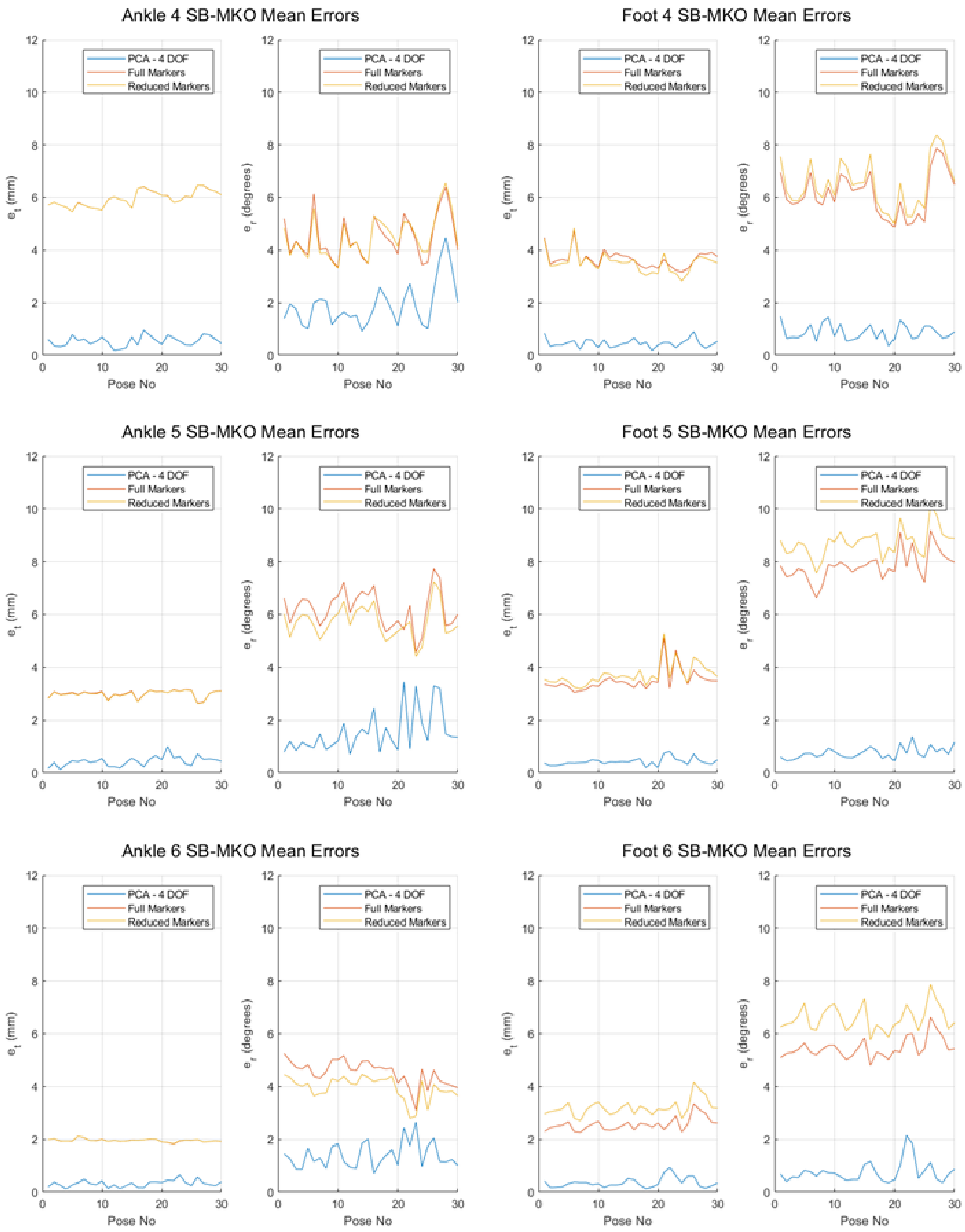


*Figure S 4: Translational and rotational error for the Ankle and the Foot of each subject and posture: blue line represent the error associated with the use of four individual subject as a reference; red line di error associated with SB-MKO using the whole foot ten marker set; the yellow line the error associated with SB-MKO using only four marker on the foot.*

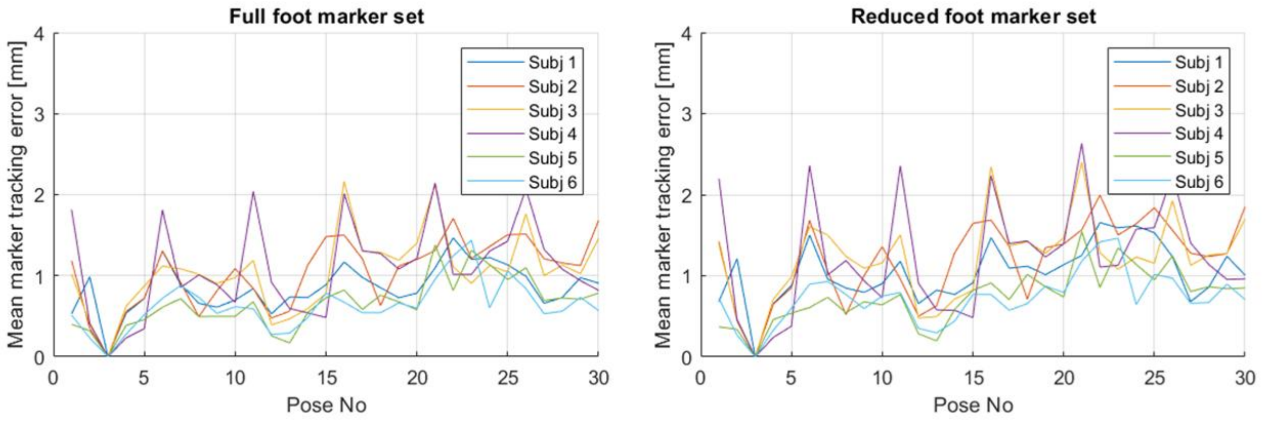


*Figure S 5: Marker residual error after SB-MKO applied to each subject using full (left) or reduced (right) foot marker set*
